# High-throughput allelic expression imbalance analyses identify candidate breast cancer risk genes

**DOI:** 10.1101/521013

**Authors:** Mahdi Moradi Marjaneh, Haran Sivakumaran, Kristine M Hillman, Susanne Kaufmann, Nehal Hussein, Luize G Lima, Sunyoung Ham, Siddhartha Kar, Jonathan Beesley, Laura Fachal, Douglas F Easton, Alison M Dunning, Andreas Möller, Georgia Chenevix-Trench, Stacey L Edwards, Juliet D French

## Abstract

Fine-mapping of breast cancer GWAS regions has identified 195 high confidence signals containing more than 5,000 credible causal variants (CCVs). The CCVs are predominantly noncoding and enriched in regulatory elements and thus may confer the risk by altering gene expression in *cis*. We analyzed allelic expression imbalance (AEI) of genes surrounding known breast cancer signals, using normal breast and breast tumor transcriptome data and imputed genotypes. Fourteen genes, including *NTN4*, were identified whose expression was associated with CCV genotype. We showed that CCVs at this signal were located within an enhancer that physically interacts with the *NTN4* promoter. Furthermore, knockdown of *NTN4* in breast cells increased cell proliferation *in vitro* and tumor growth *in vivo*. Here, we present the most comprehensive AEI analysis of breast cancer CCVs resulting in identification of new candidate risk genes.

## INTRODUCTION

The influence of common genetic variation on gene expression underlies a considerable proportion of the heritability associated with complex traits. Several studies indicate that the majority of trait-associated variants identified by Genome Wide Association Studies (GWAS) act by modulating gene expression in *cis* through altered distal and proximal regulatory elements^1^. Mapping of expression quantitative trait loci (eQTL), where genetic variants are tested for association with gene expression, is widely used to identify genes that are regulated by trait-associated variants. Several studies have shown that eQTLs are enriched in cell types relevant to the trait of interest^2,3^. For example, T cell-specific eQTLs are over-represented for autoimmune risk alleles and monocyte-specific eQTLs for Alzheimer’s and Parkinson’s disease alleles^3^. These studies highlight the importance of using relevant tissue or cell types for functional studies.

An alternative method, allelic expression imbalance (AEI) analyses, can identify associations between an imbalance of allelic expression and trait-associated variants. An advantage of AEI analyses is that associations between gene expression and genotype can be detected using substantially fewer samples as AEI is measured within individuals, thereby controlling for cellular environment and *trans*-acting genetic factors. A handful of studies have performed AEI analyses to identify genes whose expression is associated with nearby GWAS variants. For example, we and others have identified associations between AEI and risk variants for breast cancer (*ESR1, COX11* and *HELQ*)^4-6^, follicular lymphoma (*HLA-DQB1*)^7^ and high-density lipoprotein-cholesterol levels (*MMAB*)^8^.

GWAS, together with fine-mapping, have identified 196 genetic signals (with conditional p-values<10^−6^) associated with breast cancer risk^9^. However, identifying the credible causal variants (CCVs) driving the association and the genes affected by those variants is challenging. A recent study by Fachal et al has defined the CCVs as those with p-values within two orders of magnitude of the most significant CCV within the signal^9^. A signal at 17q21.31 containing 2277 CCVs co-segregates with a previously described large genomic rearrangement^10^. Following exclusion of the 17q21.31 signal, Fachal et al identified 5117 CCVs across 195 breast cancer risk signals. The majority of the CCVs fall within noncoding sequences and, in particular, they are enriched in genomic regions marked for regulatory activity suggesting that many of them may act by modulating gene expression in *cis*. In this study, we performed AEI analyses of genes surrounding the 195 breast cancer GWAS signals and identified 14 genes whose AEI was associated with CCVs (q-value<0.01). For one gene, *NTN4*, we provide mechanistic insight into how breast cancer CCVs influence *NTN4* expression, breast cancer cell proliferation and tumor development.

## RESULTS

### AEI analyses identified 14 candidate breast cancer risk genes

To identify new breast cancer risk genes whose expression is associated with breast cancer CCVs we used GTEx and TCGA breast samples with available RNA-seq and imputation data, including 81 GTEx normal, 46 TCGA normal, and 669 TCGA tumor samples. We were able to retrieve imputed genotypes for 4441 CCVs spanning over 190 signals. For each individual, from the genes within 1Mb up- and downstream of the CCVs, we selected those with at least one heterozygote transcribed single nucleotide polymorphism (htSNP), including 2195 and 2068 genes in the normal and tumor datasets, respectively. The major transcribed allele fraction (MTAF) representing AEI was then computed on each htSNP site and averaged across each gene. On average ∼2 htSNPs were available to calculate AEI per gene (Supplementary Fig. 1a).

To assess overall AEI distribution, we pooled AEI values from all genes. We observed a clear positive skew in normal (2.3) and tumor (1.8) samples, and a bimodal pattern with a main peak at AEI of 0.55 and a small but distinct peak at AEI of 1, indicating the majority of gene-sample pairs show a limited deviation from allelic expression balance while a small proportion represent extreme AEI (Supplementary Figs 1b,c). The samples were then clustered into homozygous and heterozygous classes for each CCV to test the association between AEIs and CCV genotypes. Using data from the normal samples, we were able to test 705 genes for association with CCV genotypes, representing 820 testable signal-gene pairs. The minimum p-value from CCVs within each signal was used to represent association between that signal and the corresponding gene. In general, we observed stronger association between signals and corresponding genes when they were located closer in the genome (*P*=7.7e-03; Supplementary Fig. 1d), consistent with regulatory variants more commonly acting in *cis.* Overall, in the normal breast samples, 133 genes showed evidence of AEI in heterozygous individuals (*P*<0.05; **Supplementary Table 1**), of which 14 genes passed multiple testing correction (q-value<0.01; Fig. 1 and Table 1). Conversely, analysis of the breast tumor samples resulted in 753 testable signal-gene pairs (651 genes) including 84 genes with evidence of AEI in heterozygotes (*P*<0.05; **Supplementary Table 2**), of which two genes passed multiple testing correction (q-value<0.01; Fig. 1 and Table 1).

**Table 1.** Candidate breast cancer risk genes identified by AEI analyses.

| Dataset | Gene name | CCV rsID | No. heterozygotes<br>(AEI mean) | No. homozygotes<br>(AEI mean) | p-value | q-value |
| --- | --- | --- | --- | --- | --- | --- |
| normal | *LGR6* | rs4245706 | 41 (0.64) | 34 (0.57) | 1.65E-06 | 7.06E-05 |
| normal | *ZC3H11A* | rs11240552 | 51 (0.58) | 37 (0.54) | 4.52E-06 | 1.40E-04 |
| normal | *BARX2* | rs12787230 | 25 (0.62) | 28 (0.57) | 4.28E-05 | 9.03E-04 |
| normal | *GATAD2A* | rs8101499 | 61 (0.58) | 11 (0.54) | 1.11E-04 | 1.47E-03 |
| normal | *CASP8* | rs3769821 | 53 (0.62) | 44 (0.58) | 4.27E-04 | 3.53E-03 |
| normal | *PPFIA1* | rs78540526 | 5 (0.64) | 63 (0.57) | 6.84E-04 | 5.08E-03 |
| normal | *CRLF3* | rs6505216 | 10 (0.66) | 16 (0.6) | 7.79E-04 | 5.08E-03 |
| normal | *NTN4* | rs17356907 | 32 (0.59) | 60 (0.55) | 1.17E-03 | 7.28E-03 |
| normal | *HSPA4* | rs61246276 | 17 (0.59) | 9 (0.53) | 1.37E-03 | 8.06E-03 |
| normal | *ZNF500* | rs576214 | 30 (0.63) | 38 (0.59) | 1.84E-03 | 9.27E-03 |
| normal | *TMEM80* | rs7942159 | 35 (0.67) | 15 (0.58) | 1.87E-03 | 9.27E-03 |
| normal | *RAD23B* | rs60037937 | 15 (0.57) | 22 (0.52) | 2.01E-03 | 9.30E-03 |
| normal | *STK38L* | rs61920241 | 20 (0.6) | 39 (0.55) | 2.03E-03 | 9.30E-03 |
| tumor | *KLHDC7A* | rs3007733 | 85 (0.84) | 55 (0.72) | 8.00E-06 | 8.77E-04 |

**Figure 1.**
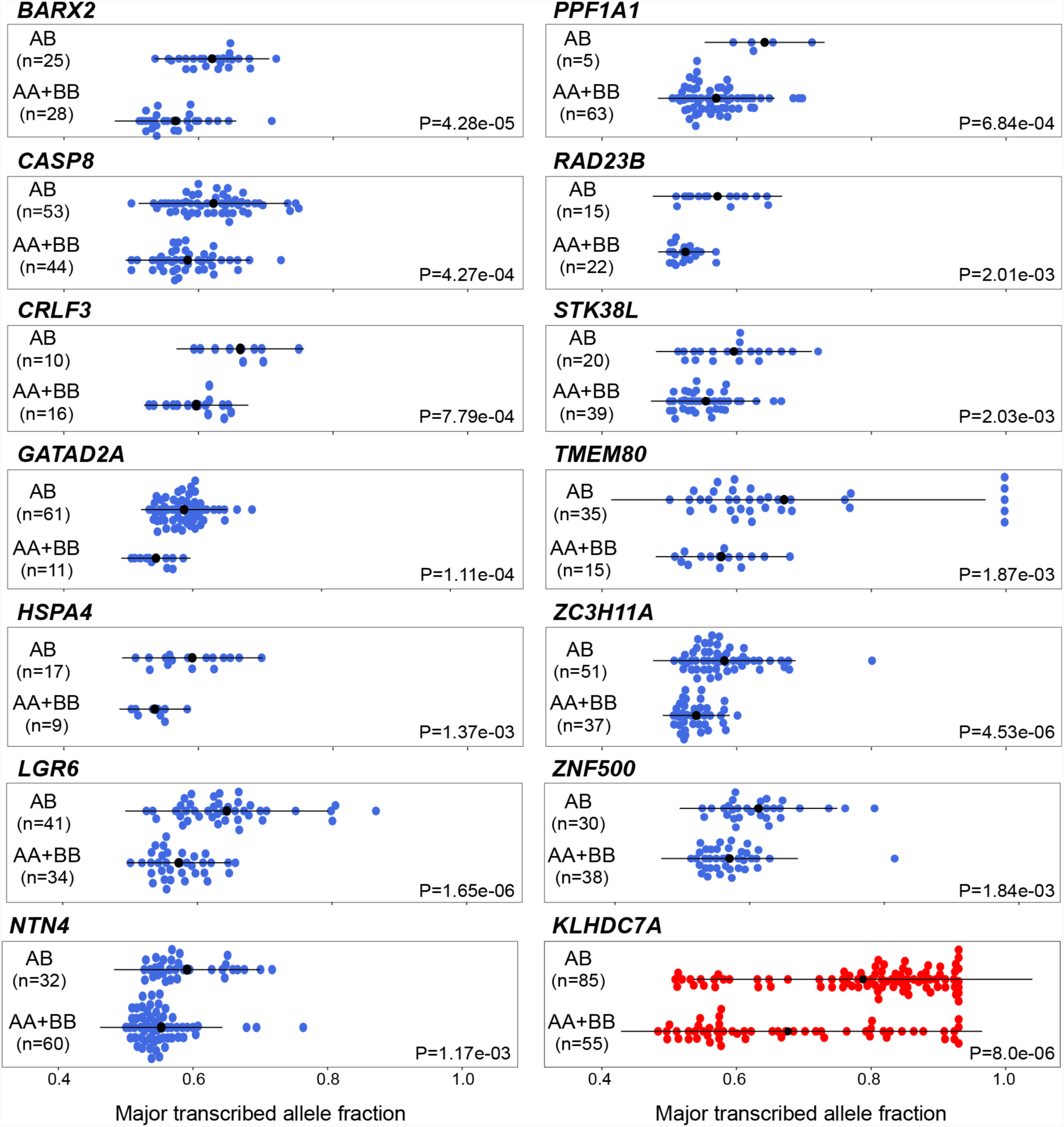
AEI by genotypic status at breast cancer CCVs. For each gene, samples are classified according to the genotypes at the corresponding CCV; heterozygous (AB) versus homozygous (AA+BB). Blue dots indicate the average major transcribed allele fraction of htSNPs across each gene representing AEI for that gene in normal breast samples from TCGA or GTEx breast datasets. Red dots indicate corresponding measurements from TCGA breast tumor datasets. Black dots and whiskers represent means ± 1 SD. *P*-values were calculated by comparing AEIs between heterozygous and homozygous groups using a two-tailed t-test adjusted for variance.

Each of the 16 genes identified by the AEI analyses had multiple tested htSNPs. Therefore, we assessed whether their AEI measurements were consistent across intra-htSNPs. Fourteen genes (*BARX2, CASP8, CRLF3, GATAD2A, HSPA4, KLHDC7A, LGR6, NTN4, PPFIA1, RAD23B, STK38L, TMEM80, ZC3H11A* and *ZNF500*) showed consistent AEI across individual htSNPs (Supplementary Fig. 2a,b, Supplementary Table 3). For each gene, only a small number of samples (<10%) had heterogeneous AEIs across individual htSNPs (*P*_heterogeneity_<0.05). However, for *HLA-A* and *RPS23*, the majority of samples showed heterogeneous AEIs (92% and 71%, respectively) and were therefore excluded from further analysis. It is possible that isoform-specific AEI may have contributed to the heterogeneity of AEI measurements across these two genes.

### Breast cancer CCVs distally regulate the AEI gene, *NTN4*

At six AEI-identified genes (*CASP8, GATAD2A, HSPA4, KLHDC7A, LGR6* and *ZC3H11A*), breast cancer CCVs are located in the promoter regions, suggesting the CCVs may confer allelic imbalance through altered promoter activity. Moreover, for six genes (*BARX2, GATAD2A, HSPA4, LGR6, NTN4* and *ZC3H11A*), our recent capture Hi-C data has indicated that chromatin looping occurs between a region containing the breast cancer CCVs and the promoter of the corresponding gene^11^. Notably, for *GATAD2A, HSPA4* and *ZC3H11A*, there are CCVs in the promoter and in distal interacting regions, and therefore it would be difficult to determine the CCVs responsible for the observed AEI. One example of a distal CCV associated with AEI is at chromosome 12q22, where genetic fine-mapping identified one risk signal that contains two CCVs (rs61938093 and rs17356097). Both CCVs fall within a putative regulatory element (PRE), marked by open chromatin, which frequently participates in long-range chromatin interactions with the *NTN4* promoter region in B80T5 and MCF10A nontumorigenic breast cell lines (Fig. 2a). Silencing of the PRE by targeting a nuclease-defective dCas9 fused to the Kruppel-associated box (dCas9-KRAB) reduced *NTN4* expression in B80hTERT1 cells, suggesting that the PRE acts as an enhancer (Fig. 2b). Luciferase reporter assays further confirmed strong enhancer activity of the PRE on the *NTN4* promoter. However, inclusion of the risk-associated CCV alleles did not alter enhancer activity (Fig. 2c). Notably, in T47D, a breast cancer cell line heterozygous for the CCVs, allele-specific 3C showed a preference for the protective allele (Figs 2d,e, Supplementary Fig. 3a,b), suggesting that risk alleles may abrogate looping between the enhancer and *NTN4* which in turn may reduce *NTN4* expression. Using electromobility shift assays (EMSAs) we showed that rs61938093 altered protein binding *in vitro*, suggesting this is the likely functional variant at the signal (Fig. 2f). However, we were unable to identify the specific protein that binds the risk allele.

**Figure 2.**
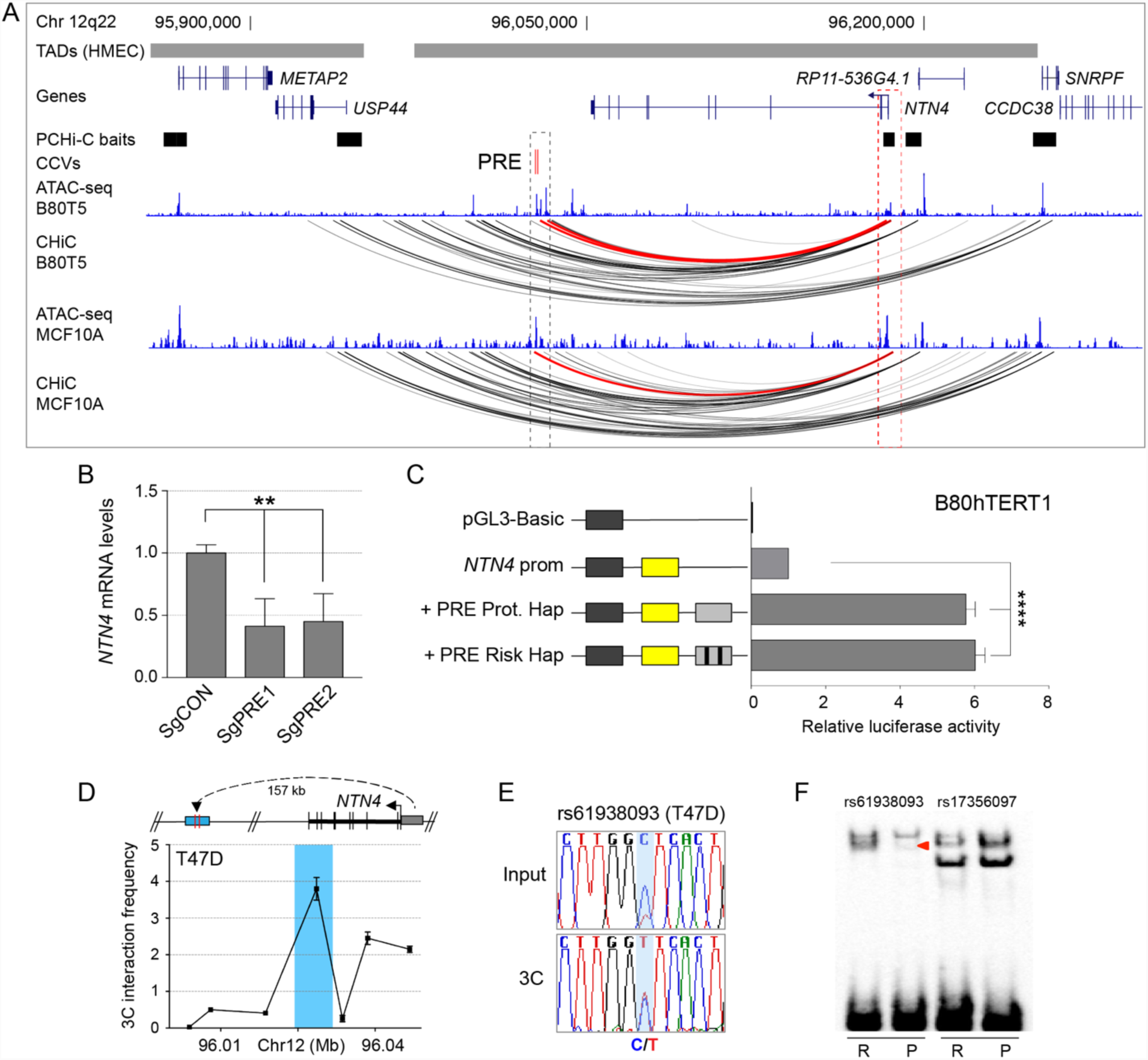
Breast cancer CCVs distally regulate *NTN4*. (**a**) WashU genome browser showing topologically associating domains (TADs) as horizontal gray bars above GENCODE annotated coding genes (blue). The PCHi-C baits are depicted as a black boxes. The putative regulatory element (PRE) containing the CCVs is shown as red colored vertical lines. The ATAC-seq tracks for B80T5 and MCF10A breast cells are shown as blue histograms. Capture Hi-C (CHi-C) chromatin interactions are shown as. Red arcs depict chromatin looping between CCVs and the *NTN4* promoter region. (**b**) dCAS9-KRAB was targeted to the PRE using two different sgRNAs (sgPRE1 and sgPRE2) in B80hTERT1 breast cells. SgCON contains a non-targeting control guide RNA. Gene expression was measured by qPCR and normalized to *beta-glucuronidase* (GUSB) expression. Error bars, SEM (n=3). *P*-values were determined by two-way ANOVA followed by Dunnett’s multiple comparisons test (* * p<0.01). (**c**) Luciferase reporter assays following transient transfection of B80hTERT1 breast cells. The PRE containing either the protective (Prot.) or risk alleles of rs61938093 and rs1735097 was cloned into *NTN4*-promoter driven luciferase constructs. Error bars, SEM (n=3). *P*-values were determined by two-way ANOVA followed by Dunnett’s multiple comparisons test (* * * * p<0.0001). (**d**) 3C interaction profiles between the *NTN4* promoter and the genomic region containing the PRE in T47D 3C libraries generated with HindIII. A physical map of the region interrogated by 3C is shown (top panel), with the blue shading representing the position of the PRE and the anchor point set at the *NTN4* promoter. Representative 3C profile is shown (bottom panel). Error bars, SD (n=3). (**e**) Sequencing chromatogram of T47D input versus the 3C PCR product that showed allele specific looping at rs61938093. One of three independent 3C libraries is shown. (**f**) EMSA for oligonucleotide duplexes containing SNPs rs61938093 or rs1735097 with either the risk allele (R) or protective allele (P) as indicated, assayed using B80hTERT1 nuclear extracts. Arrowhead indicates band mobility differences between alleles.

### *NTN4* knockdown increases cell proliferation *in vitro* and tumor growth in mice

We examined expression of *NTN4* in normal and cancerous breast tissues using TCGA RNA-seq data. *NTN4* was more highly expressed in normal tissue compared to adjacent tumor samples (Fig. 3a) and expressed across the histological subtypes, albeit with lower expression in the basal subtype (Fig. 3b). To assess the effect of reduced *NTN4* on cell proliferation, we knocked down *NTN4* using siRNA and showed that reduced *NTN4* significantly increased cell proliferation (Fig. 3c, Supplementary Fig. 3c). To assess the effect of reduced *NTN4* on tumor growth, we stably depleted *NTN4* in MCF7 cells by targeting dCAS9-KRAB to the promoter of *NTN4*, and injected the cells in the mammary fat pad of nude mice. Compared to control MCF7 cells containing non-targeting sgRNA, *NTN4* depletion led to a marked increase in tumor growth (Figs 3d,e, Supplementary Fig. 3d), which was reflected in increased tumor weight (Fig. 3f).

**Figure 3.**
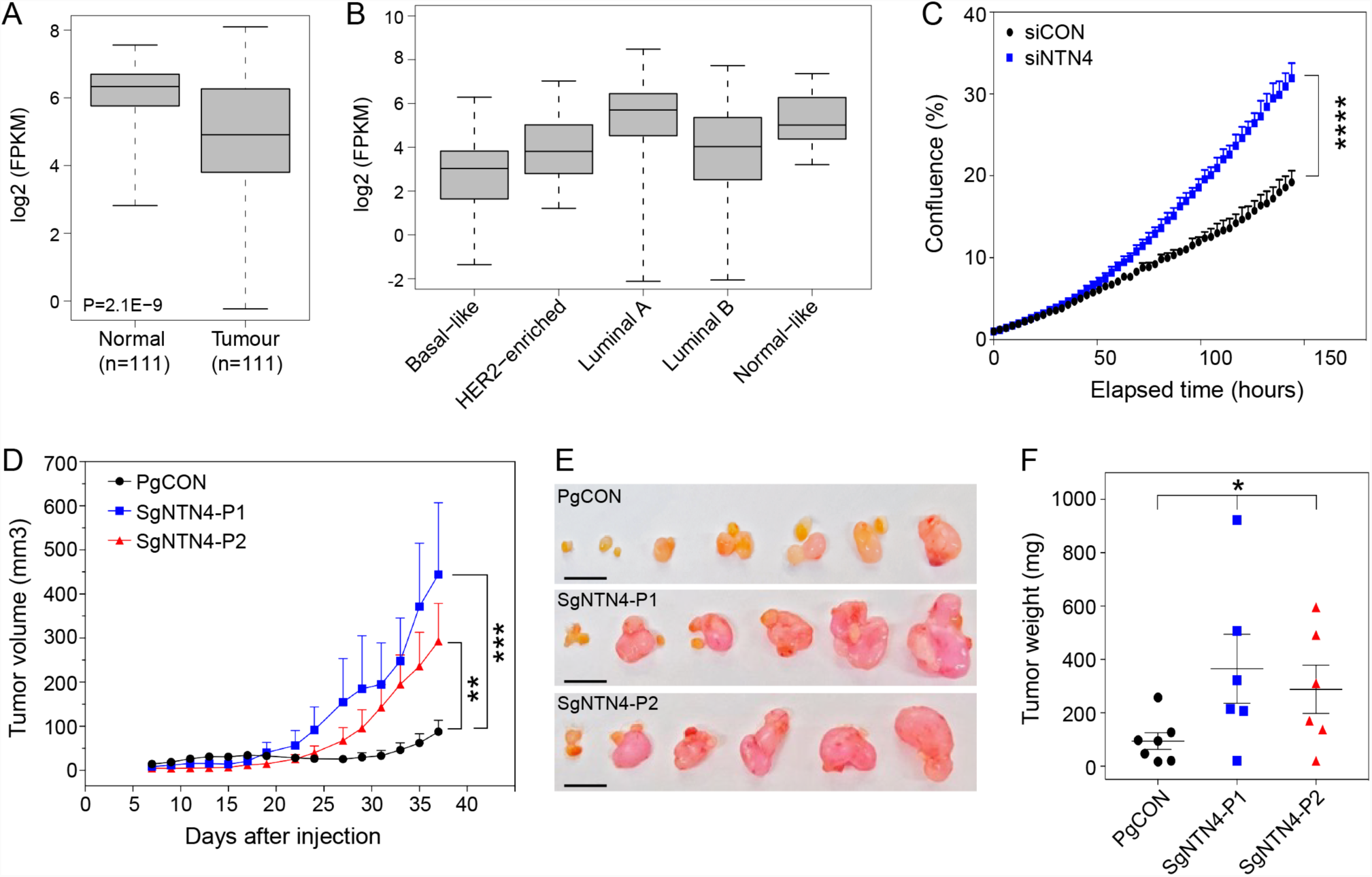
*NTN4* depletion promotes cell proliferation and tumor formation. (**a**) *NTN4* expression in normal breast and paired tumor tissue samples from TCGA. *P*-values were determined using a two-tailed t-test. (**b**) *NTN4* expression in breast tumors stratified by PAM50 molecular subtypes (n=841). (**c**) Proliferation of MCF7 breast tumor cells transfected with a control (siCON) or *NTN4* (siNTN4) Smartpool siRNA. Cells were grown in 24-well plates and confluency of the wells was measured by the IncuCyte live-cell imaging system. Results represent relative cell growth rates. Error bars, SEM (n=4). (**d**) MCF7-control (SgCON) or MCF7-dCas9-KRAB NTN4 repressed cells (SgNTN4-P1/P2) were orthotopically injected into the mammary fat pads of nude mice. Tumor growth curves for each group are shown. Values are shown as average tumor volumes at each time point. Error bars, SEM (n=6-7 mice per group). (**e**) Tumors of individual mice were dissected at day 38 post-injection. The scale bar represents 1 cm. (**f**) Plot of the individual weights of tumors with mean and SEM shown by cross-bar and error bars. (**d, f**) Mann-Whitney *U* test was used to compare differences between groups (* p<0.05, * * p<0.01, * * * p<0.001).

## DISCUSSION

Identifying the target genes of GWAS variants is challenging and requires multiple bioinformatic and functional approaches. In this study, we used AEI analyses to link breast cancer CCVs to their likely target genes. Using RNA-seq data and imputed genotypes from two breast datasets (TCGA and GTEx), we identified 14 genes in which AEI was associated with breast cancer CCVs at a strict multiple testing threshold. Thirteen genes were identified from 127 normal breast samples and one gene from 669 breast tumor samples. The low number of genes identified from the tumor dataset likely reflects the heterogeneous nature of tumor samples and/or high levels of AEI caused by cancer related mechanisms such as copy number variation, aberrant DNA methylation or acquired regulatory mutations^12^.

We used MTAF as a measure of AEI for each htSNP. This allowed us to average AEIs over multi-htSNP genes (more than 50% of the tested genes) and reduced the influence of sequencing errors at individual htSNPs. Furthermore, we could pool two classes of homozygous genotypes, providing greater statistical power when comparing to heterozygotes. We also retrieved SNP genotypes from high-confidence SNP array calls imputed to the 1000 Genome reference panel and therefore did not rely on setting a minimum cut-off for MTAF when deriving genotypes from RNA-seq data. However, while powerful, we acknowledge this AEI-based approach has some limitations. The multi-level criteria we applied including the minimum expression level or stringent significance threshold may have compromised sensitivity when genes are lowly expressed or only weakly regulated by CCVs. False positives may also arise from htSNP-related mapping biases, caused by repeat sequences or low-complexity regions. In addition, sequencing reads containing the reference allele at a htSNP site are more likely to be mapped, which would result in a false AEI^13,14^.

Five of the genes we identified have previously been identified through eQTL studies using either breast tumor or normal breast tissue, namely *GATAD2A, CASP8, NTN4, HSPA4* and *RPS23*^*15*^, while *KLHDC7A* was previously identified through a transcription wide association study using expression data from breast tissue^16^. In addition, several AEI-identified genes have previously been implicated in cancer biology. *CASP8* encodes an apoptotic enzyme that functions as an initiator caspase in the extrinsic apoptotic signalling pathway^17^. CASP8 deficiency can contribute to cancer development through multiple processes, including effects on cellular transformation, adhesion and migration^18,19^. *BARX2* encodes a homeobox transcription factor that can function as a tumor suppressor and its down regulation is a predictor of poor prognosis in different cancer types^20-22^. BARX2 can bind the estrogen receptor (the target of tamoxifen) and influence estrogen-dependent growth and cellular invasion^23^. Overexpression of *PPFIA1* has also been reported as a potential predictor of metastatic relapse and poor prognosis in estrogen receptor positive/nodal negative breast tumors^24^. *LGR6* is reported to promote cancer development and, when suppressed, induces apoptosis^25^. Moreover, the protein encoded by *LGR6* has been specifically detected on the surface of progenitor cells that likely give rise to luminal-type mammary tumors^26^.

Of particular interest was *NTN4*, which encodes the Netrin-4 secreted protein, implicated in various developmental processes including axon guidance, angiogenesis, and mammary and lung morphogenesis^27^. Several studies have implicated *NTN4* in breast cancer progression. For example, reduced *NTN4* is reported to promote proliferation, migration and invasion of breast cancer cells by promoting epithelial to mesenchymal transition^28^. In addition, NTN4 has been shown to be an independent biomarker for prognosis of survival in breast cancer ^29,30^. We and others have demonstrated that SNPs can alter chromatin loop formation between promoters and enhancers^31,32^. Here, we provide evidence that the same mechanism may explain how breast cancer CCVs alter *NTN4* expression and that suppressed *NTN4* increases cancer-related processes including cell proliferation and tumor growth.

In summary, we present the most comprehensive AEI analysis linking breast cancer CCVs to their target genes. Fourteen genes were identified, including some potential cancer driver genes, but many with no reported role in breast cancer biology. Future work will be required to confirm the role of these genes in breast cancer development, which could ultimately lead to new avenues for breast cancer prevention or therapy.

## METHODS

### Allelic expression imbalance (AEI) analyses

CCVs associated with breast cancer risk were tested for association with AEI of genes within 1Mb up- and downstream using RNA-seq data and imputed genotypes from The Cancer Genome Atlas (TCGA; https://portal.gdc.cancer.gov/) and The Genotype-Tissue Expression (GTEx; https://dbgap.ncbi.nlm.nih.gov/) breast datasets. TCGA imputation was performed for individuals with European ancestry (N=701) using 1000 Genomes version 5 reference panel and without pre-phasing. From dbSNP human Build 146 we extracted SNPs located within exonic regions of the target genes and computed RNA-seq read counts on SNP sites for reference and alternative alleles using *bam-readcount* (https://github.com/genome/bam-readcount). Sequencing reads with base quality < 15 at a position were not considered for allele counting. For each individual and each gene, heterozygote transcribed SNPs (htSNPs) were identified using imputed genotypes and RNA-seq read counts (>15x RNA-seq read depth). For each gene, AEI of the htSNPs were computed and averaged to represent AEI for that gene (ranging from 0.5 to 1). Homogeneity of AEI across individual htSNPs within a gene was assessed using the MBASED R package. For the analysis of tumor samples, those with copy number alteration for a given gene were excluded. For each CCV, samples were classified into homozygous and heterozygous groups. CCV-gene pairs with less than five samples in either the homozygous or heterozygous group as well as those with a higher AEI mean in the homozygous group were excluded. AEI was compared between heterozygous (AB) and homozygous (AA+BB) samples using a two-sided t-test adjusted for variance. *P* significance level was adjusted for multiple testing using the q-value method.

### Cell lines

Breast cancer cell lines T47D and MCF7 were maintained in RPMI 1640 medium supplemented with 10% (v/v) fetal bovine serum, 10 μg/ml insulin and 1% (v/v) antibiotic-antimycotic (Life Technologies). B80hTERT1 normal breast epithelial cells (Roger Reddel, Children’s Medical Research Institute, Sydney, Australia) were maintained in 1:1 MCDB 170 and RPMI 1640 media supplemented with 10% (v/v) fetal bovine serum and 1% (v/v) antibiotic-antimycotic. Cells were cultured in a humidified 5% CO_2_ atmosphere at 37°C. Cell-line quality control included short tandem repeat profiling and *Mycoplasma* contamination screening.

### Chromosome conformation capture (3C)

3C libraries were generated using HindIII from T47D cells as previously described^33^. 3C interactions were quantified by qPCR using the Rotor-Gene 6000 platform, annealing at 66°C for 30 secs in the presence of 25 μM Syto9 fluorescent dye (Life Technologies). BAC clones were used to create artificial libraries of ligation products to normalize for PCR efficiency. Data were normalized to the signal from the BAC clone library and between replicates by reference to a region within *GAPDH*. 3C primers are listed in **Supplementary Table 4**.

### Allele specific 3C

T47D 3C libraries or T47D genomic DNA were amplified with allele specific PCR primers (**Supplementary Table 4**). PCR products were electrophoresed through 2% (w/v) agarose and bands excised for DNA extraction (QIAGEN). Purified amplicons were Sanger sequenced by the Australian Genome Research Facility (AGRF).

### Luciferase reporter assays

The *NTN4* promoter-driven luciferase reporter construct was generated by inserting a PCR amplified genomic fragment into the KpnI/HindIII sites of the pGL3-basic vector (Promega). A 1999bp fragment containing the PRE, with either the risk or protective alleles, were synthesized as gBlocks (Integrated DNA Technologies) and then cloned into the BamHI/SalI sites of the *NTN4*-promoter construct (genomic coordinates and primers are listed in **Supplementary Table 4**). B80hTERT1 cells were transfected with the reporter constructs and a control pRL-TK *Renilla* plasmid using Lipofectamine 3000 (Life Technologies). Luciferase activity was measured 24 h post-transfection using the Dual-Glo Luciferase System (Promega). To correct for any differences in transfection efficiency, *Firefly* luciferase activity was normalized to *Renilla* luciferase and the results expressed relative to the *NTN4*-promoter construct, which had a defined activity of 1. *P*-values were determined by two-way ANOVA followed by Dunnett’s multiple comparisons test.

### Electromobility shift assays

Nuclear lysates from B80hTERT1 cells were prepared using the NE-PER nuclear and cytoplasmic protein extraction kit (Thermo Fisher). Biotinylated oligonucleotides representing either the risk or protective allele were synthesized (Integrated DNA Technologies; **Supplementary Table 4**) and annealed to form double-stranded duplexes. Nuclear lysates (5 μg) and duplexes (10 fmol) were combined in binding reactions containing 10% (v/v) glycerol, 1 mM DTT, 0.5 μg poly(dI:dC) (Sigma-Aldrich), protease inhibitors (Roche), and 20 mM HEPES (pH 7.4) at 25°C for 15 min. Reactions were resolved by electrophoresis in 10% (w/v) Tris-borate-EDTA polyacrylamide (Lonza) and transferred to positively-charged nylon membranes by semi-dry transfer (Bio-Rad). Membranes were assayed for gel shift complexes using the LightShift Chemiluminescent EMSA kit (Thermo Fisher) and visualized with the XX6 gel documentation system (Syngene).

### CRISPR interference (CRISPRi)

CRISPRi was performed with the lentiviral vector pHR-SFFV-dCas9-BFP-KRAB (dCas9-KRAB; a gift from Stanley Qi & Jonathan Weissman, Addgene plasmid #46911). Single-guide RNAs (sgRNAs) targeting the *NTN4* promoter or PRE were designed (**Supplementary Table 4**) and synthesized (Integrated DNA Technologies) for cloning into the lentiviral vector pgRNA-humanized (a gift from Stanley Qi, Addgene plasmid #44248). Lentiviral particles were produced from HEK293 cells transfected with accessory plasmids pCMV-dR8.91 and pCMV-VSV-G (gifts from David Harrich, QIMR Berghofer), along with either dCas9-KRAB or a pgRNA construct, using Lipofectamine 2000 (Life Technologies). Supernatants from dCas9-KRAB and pgRNA cultures were mixed and transduced into MCF7 or B80hTERT1 target cells. Transductants expressing both dCas9-KRAB (co-expressing blue fluorescent protein) and pgRNA (co-expressing mCherry) were enriched by FACS on the Aria IIIu platform (Becton-Dickinson). Knockdown of *NTN4* expression was confirmed by TaqMan qPCR gene expression assays (Life Technologies).

### RNA interference (RNAi)

MCF7 cells were transfected with either ON-TARGETplus negative control or *NTN4* siRNA Smartpools (Dharmacon; **Supplementary Table 4**) using RNAiMAX transfection reagent (Life Technologies) at a concentration of 5 pmol per 10^6^ cells. Knockdown of *NTN4* was confirmed by TaqMan qPCR gene expression assay 72 h post-transfection.

### Cell proliferation assay

The IncuCyte live cell imaging platform (Essen Bioscience) was used to measure cell proliferation by time course. Treated cells were plated at low confluency and imaged at 10× magnification every 3 h over 7 d. The rate of change of cell confluency was determined using ZOOM 2016A (Essen Bioscience) and Prism (GraphPad) software and compared to control treated cells.

### Mouse xenografts

Female BALB/c-Foxn1^nu^/Arc mice were subcutaneously implanted with 17β-estradiol (720 μg, 90 d release; Innovative Research of America) at 8 weeks. MCF7 control-CRISPRi or *NTN4*-CRISPRi cells were orthotopically injected into mammary fatpads 3 days later at 10^7^ cells per mouse, 6-7 mice per cell line. Tumor volumes were measured every 2 days until experimental end, at which point mice were euthanized and their tumors excised and weighed. All animal procedures were conducted in accordance with Australian National Health and Medical Research regulations on the use and care of experimental animals, and approved by the QIMR Berghofer Medical Research Institute Animal Ethics Committee (P1499).

## ACKNOWLEDGEMENTS

This work was supported by a grant from the National Health and Medical Research Council of Australia (NHMRC; 1120563). S.L.E. is an NHMRC Senior Research Fellow (1135932). G.C.T is an NHMRC Senior Principle Research Fellow (1117073). J.D.F was supported by a Fellowship from the National Breast Cancer Foundation of Australia. N.A. was co-funded by a QIMR Berghofer International PhD Scholarship and a University of Queensland Research Training Scholarship. This project has received funding from the European Union’s Horizon 2020 Marie Skłodowska-Curie Individual Fellowships programme under grant agreement No [MSCA-IF-2014-EF-656144]. The results published here are in part based upon data generated by the TCGA Research Network. The authors declare no competing financial interests.

---

**Supplementary Figure 1.**
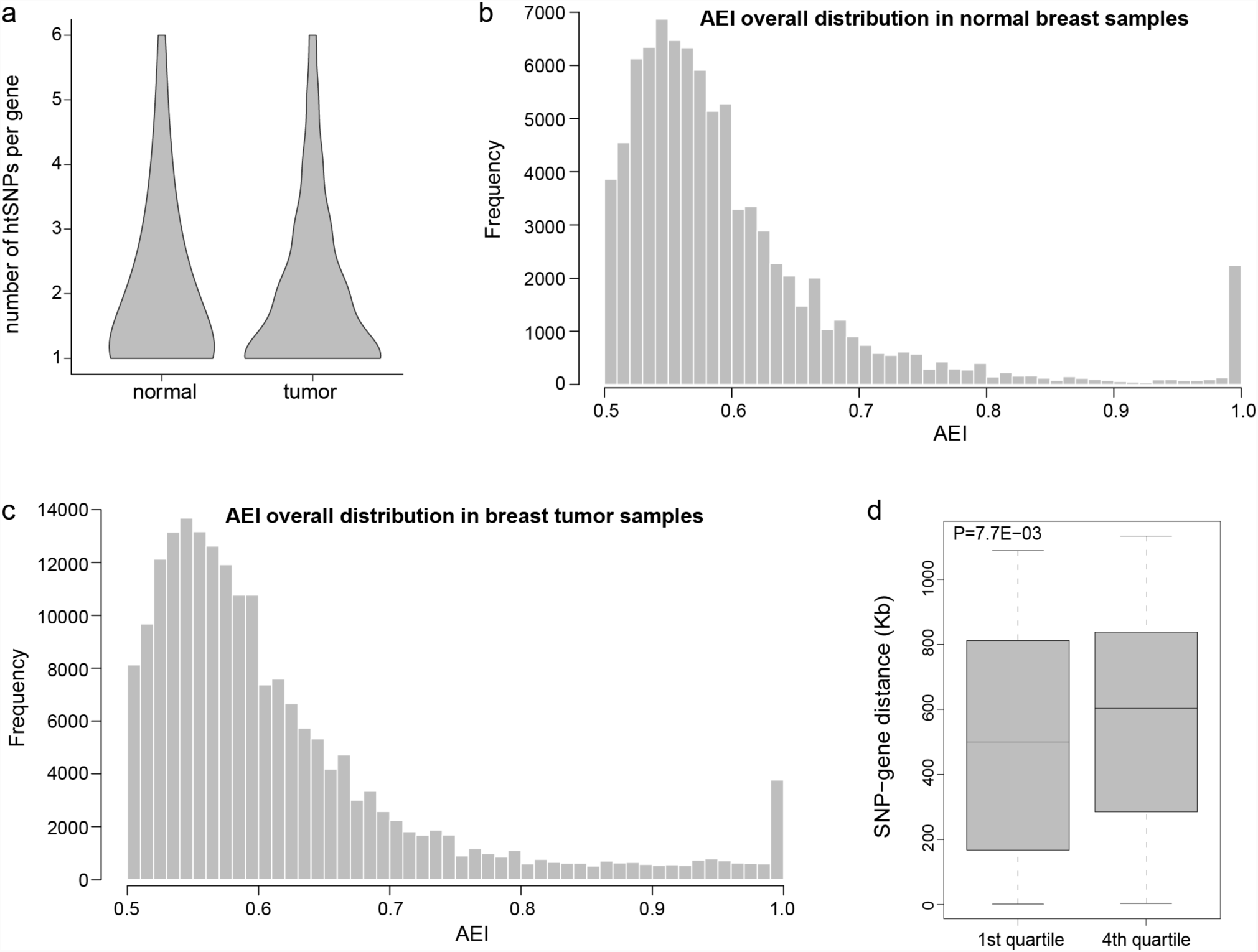
Quality control of the data used for allelic expression imbalance (AEI) analyses. (**a**) Violin plots showing overall distribution of heterozygote transcribed single nucleotide polymorphism (htSNP) counts per gene in normal breast or breast tumor samples. (**b, c**) Overall distribution of AEI measurements from genes with at least one htSNP within normal breast or breast tumor samples. (**d**) *P*-values representing the probability of association between each CCV and a nearby gene were sorted and the top and bottom quartiles compared for CCV-gene genomic distances using a two-tailed t-test.

**Supplementary Figure 2.**
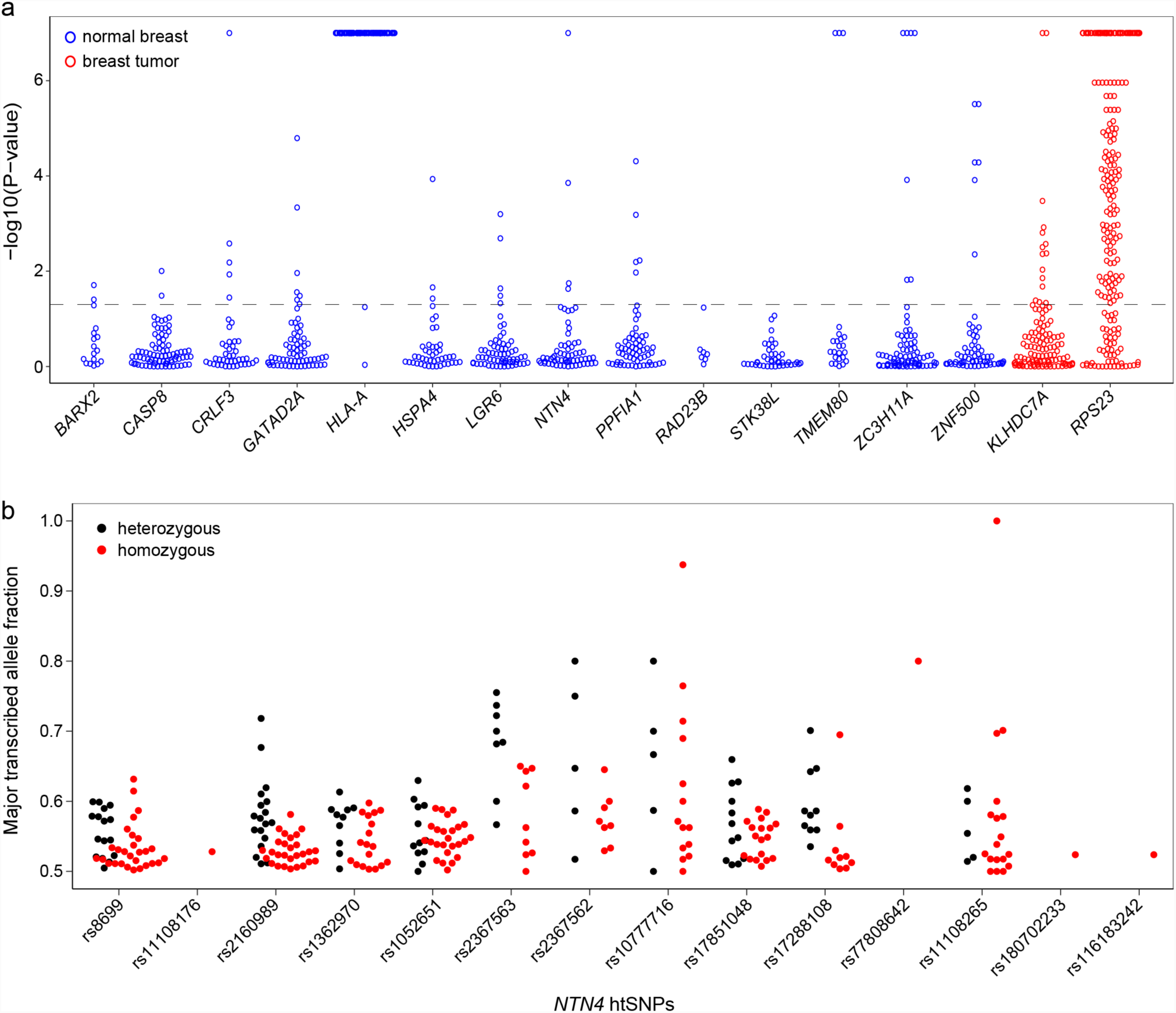
AEI consistency across htSNPs. **(a)** Each circle represents a normal breast (blue) or breast tumor (red) sample with at least two htSNPs for a corresponding gene. The y-axis shows *P*_heterogeneity_ (minus log-transformed) representing AEI variability across individual htSNPs. The black dashed horizontal line indicates level of statistical significance at *P* < 0.05. Samples above this line have a heterogeneous AEI across htSNPs for the corresponding gene. **(b)** AEI across individual *NTN4* htSNPs. Each circle represents the major transcribed allele fraction for the corresponding htSNP in samples either heterozygous (black) or homozygous (red) for the CCV.

**Supplementary Figure 3.**
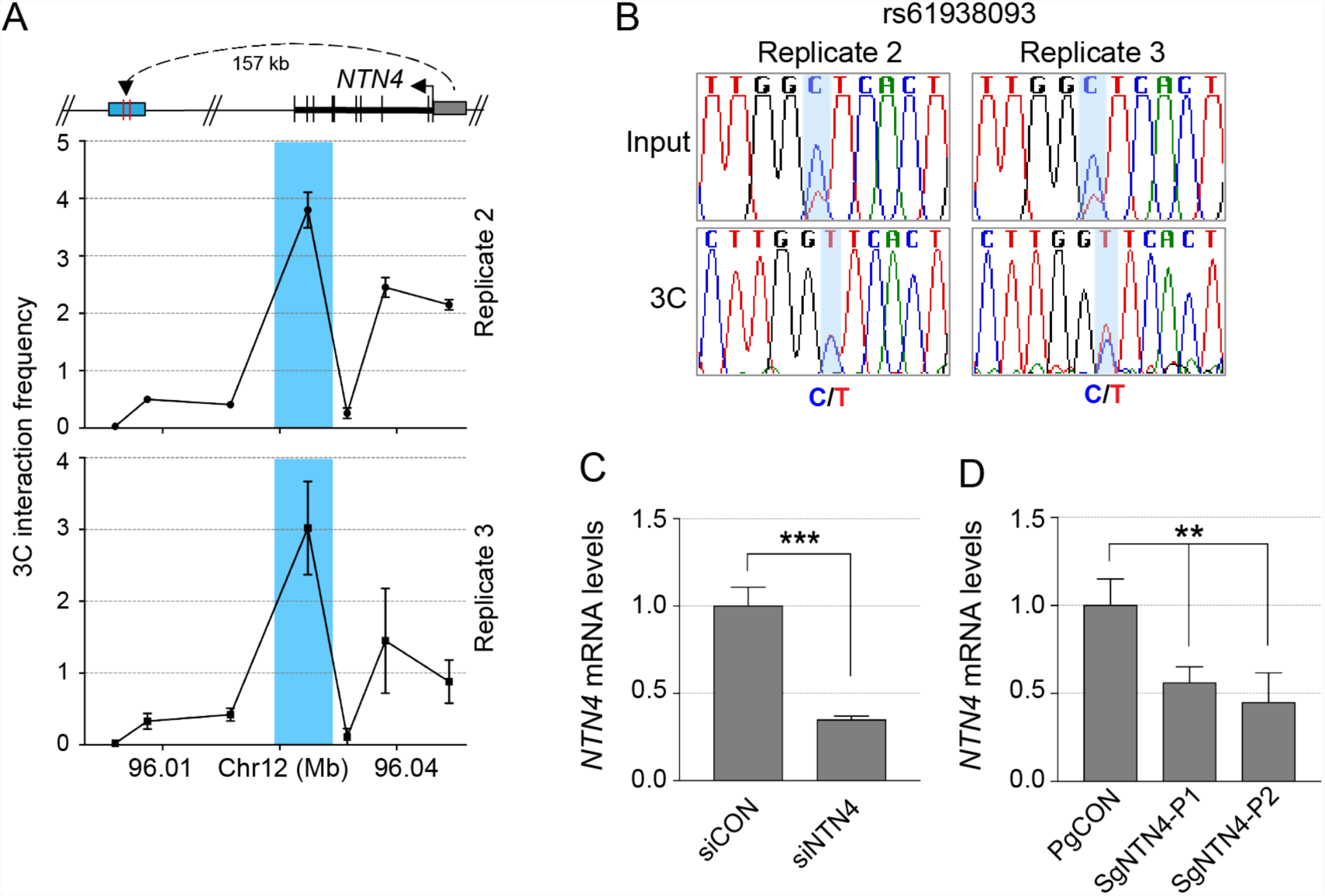
Additional in vitro data for *NTN4*. (**a**) Replicate 3C interaction profiles between the *NTN4* promoter and the genomic region containing the PRE in T47D cells. 3C libraries were generated with HindIII, with the anchor point set at the *NTN4* promoter. A physical map of the region interrogated by 3C is shown above, the blue shading represents the position of the PRE. Error bars, SD (n=3). (**b**) Replicate sequencing chromatograms of T47D input vs 3C PCR product showing allele specific looping at rs61938093. (**c**) *NTN4* depletion after transient transfection of a control (siCON) or *NTN4* (siNTN4) siRNA smartpool. *NTN4* mRNA levels were measured by qPCR and normalized to *beta-glucuronidase (GUSB)*. Error bars, SEM (n=3). *P*-values were determined with a two-tailed *t*-test (* * * p<0.001). (**d**) *NTN4* depletion in MCF7-control (PgCON) or MCF7-dCas9-KRAB *NTN4* repressed cells (SgNTN4-P1/P2). *NTN4* mRNA expression was measured by qPCR and normalized to *GUSB*. Error bars, SEM (n=3). P-values were determined by two-way ANOVA followed by Dunnetts multiple comparisons test (* * p<0.01).

## REFERENCES

1. Edwards, S.L., Beesley, J., French, J.D. & Dunning, A.M. Beyond GWASs: illuminating the dark road from association to function. Am J Hum Genet 93, 779–97 (2013).

2. Ongen, H. et al. Estimating the causal tissues for complex traits and diseases. Nat Genet 49, 1676–1683 (2017).

3. Raj, T. et al. Polarization of the effects of autoimmune and neurodegenerative risk alleles in leukocytes. Science 344, 519–23 (2014).

4. Dunning, A.M. et al. Breast cancer risk variants at 6q25 display different phenotype associations and regulate ESR1, RMND1 and CCDC170. Nat Genet 48, 374–86 (2016).

5. Darabi, H. et al. Fine scale mapping of the 17q22 breast cancer locus using dense SNPs, genotyped within the Collaborative Oncological Gene-Environment Study (COGs). Sci Rep 6, 32512 (2016).

6. Hamdi, Y. et al. Association of breast cancer risk with genetic variants showing differential allelic expression: Identification of a novel breast cancer susceptibility locus at 4q21. Oncotarget 7, 80140–80163 (2016).

7. Conde, L., Bracci, P.M., Richardson, R., Montgomery, S.B. & Skibola, C.F. Integrating GWAS and expression data for functional characterization of disease-associated SNPs: an application to follicular lymphoma. Am J Hum Genet 92, 126–30 (2013).

8. Fogarty, M.P., Xiao, R., Prokunina-Olsson, L., Scott, L.J. & Mohlke, K.L. Allelic expression imbalance at high-density lipoprotein cholesterol locus MMAB-MVK. Hum Mol Genet 19, 1921– 9 (2010).

9. Fachal, L. et al. Fine mapping of 150 breast cancer risk regions identifies 178 high confidence target genes. Submitted (2018).

10. Boettger, L.M., Handsaker, R.E., Zody, M.C. & McCarroll, S.A. Structural haplotypes and recent evolution of the human 17q21.31 region. Nat Genet 44, 881–5 (2012).

11. Beesley, J. et al. Chromatin interactome mapping at 139 independent breast cancer risk signals. Submitted (2018).

12. Mayba, O. et al. MBASED: allele-specific expression detection in cancer tissues and cell lines. Genome Biol 15, 405 (2014).

13. Heap, G.A. et al. Genome-wide analysis of allelic expression imbalance in human primary cells by high-throughput transcriptome resequencing. Hum Mol Genet 19, 122–34 (2010).

14. Castel, S.E., Levy-Moonshine, A., Mohammadi, P., Banks, E. & Lappalainen, T. Tools and best practices for data processing in allelic expression analysis. Genome Biol 16, 195 (2015).

15. Guo, X. et al. A Comprehensive cis-eQTL Analysis Revealed Target Genes in Breast Cancer Susceptibility Loci Identified in Genome-wide Association Studies. Am J Hum Genet 102, 890–903 (2018).

16. Wu, L. et al. A transcriptome-wide association study of 229,000 women identifies new candidate susceptibility genes for breast cancer. Nat Genet 50, 968–978 (2018).

17. Ashkenazi, A. Targeting the extrinsic apoptosis pathway in cancer. Cytokine Growth Factor Rev 19, 325–31 (2008).

18. Krelin, Y. et al. Caspase-8 deficiency facilitates cellular transformation in vitro. Cell Death Differ 15, 1350–5 (2008).

19. Finlay, D. & Vuori, K. Novel noncatalytic role for caspase-8 in promoting SRC-mediated adhesion and Erk signaling in neuroblastoma cells. Cancer Res 67, 11704–11 (2007).

20. Sellar, G.C. et al. BARX2 induces cadherin 6 expression and is a functional suppressor of ovarian cancer progression. Cancer Res 61, 6977–81 (2001).

21. Mi, Y. et al. Downregulation of homeobox gene Barx2 increases gastric cancer proliferation and metastasis and predicts poor patient outcomes. Oncotarget 7, 60593–60608 (2016).

22. Mi, Y. et al. Down-regulation of Barx2 predicts poor survival in colorectal cancer. Biochem Biophys Res Commun 478, 67–73 (2016).

23. Stevens, T.A. & Meech, R. BARX2 and estrogen receptor-alpha (ESR1) coordinately regulate the production of alternatively spliced ESR1 isoforms and control breast cancer cell growth and invasion. Oncogene 25, 5426–35 (2006).

24. Yang, J. et al. PPFIA1 is upregulated in liver metastasis of breast cancer and is a potential poor prognostic indicator of metastatic relapse. Tumour Biol 39, 1010428317713492 (2017).

25. Wang, F. et al. Downregulation of Lgr6 inhibits proliferation and invasion and increases apoptosis in human colorectal cancer. Int J Mol Med 42, 625–632 (2018).

26. Blaas, L. et al. Lgr6 labels a rare population of mammary gland progenitor cells that are able to originate luminal mammary tumours. Nat Cell Biol 18, 1346–1356 (2016).

27. Wilson, B.D. et al. Netrins promote developmental and therapeutic angiogenesis. Science 313, 640–4 (2006).

28. Xu, X., Yan, Q., Wang, Y. & Dong, X. NTN4 is associated with breast cancer metastasis via regulation of EMT-related biomarkers. Oncol Rep 37, 449–457 (2017).

29. Esseghir, S. et al. Identification of transmembrane proteins as potential prognostic markers and therapeutic targets in breast cancer by a screen for signal sequence encoding transcripts. J Pathol 210, 420–30 (2006).

30. Esseghir, S. et al. Identification of NTN4, TRA1, and STC2 as prognostic markers in breast cancer in a screen for signal sequence encoding proteins. Clin Cancer Res 13, 3164–73 (2007).

31. Betts, J.A. et al. Long Noncoding RNAs CUPID1 and CUPID2 Mediate Breast Cancer Risk at 11q13 by Modulating the Response to DNA Damage. Am J Hum Genet 101, 255–266 (2017).

32. Visser, M., Kayser, M. & Palstra, R.J. HERC2 rs12913832 modulates human pigmentation by attenuating chromatin-loop formation between a long-range enhancer and the OCA2 promoter. Genome Res 22, 446–55 (2012).

33. Ghoussaini, M. et al. Evidence that breast cancer risk at the 2q35 locus is mediated through IGFBP5 regulation. Nat Commun 4, 4999 (2014).

